## Supplemental figures, tables, and methods for "NRF2 activators inhibit influenza A virus replication by interfering with nucleo-cytoplasmic export of viral RNPs in an NRF2-independent manner"

**Table of contents**

**Item**	**Page**

Figure S1	2

Figure S2	3

Figure S3	4

Figure S4	5

Figure S5	6

Figure S6	7

Figure S7	8

Figure S8	8

Figure S9	9

Figure S10	10

Table S1	11

Table S2	12

Supplemental Methods	13

Figure S11	15

Figure S12	15

Figure S13	16

Figure S14	16

Figure S15	17

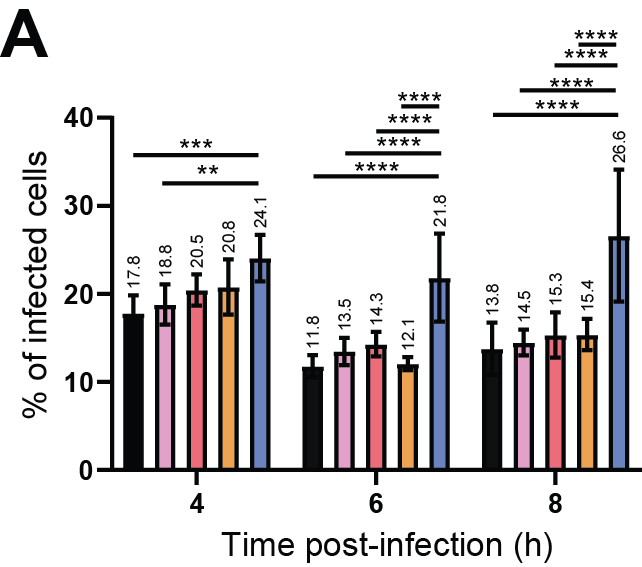

**Figure S1. Percentage of infected cells throughout an 8 h time course of IAV infection.** A549 cells were pretreated with the compounds (SEL, 1 µM; 4OI, 100 µM; BARD, 0.1 µM; SFN, 10 µM) for 12 h, were then infected with IAV PR8M (MOI=1) for 1 h and subsequently incubated in fresh medium containing the compounds. Analysis based on the same images as used for **Figure 2C-F**. Total number of cells was determined by counting DAPI-positive nuclei, and IAV infected cells by counting cells staining positive for NP in nucleus, cytoplasm or both. Data correspond to averages from 7 microscopic fields. **A.** Percentage of infected cells at 4, 6, and 8 h p.i. One-way ANOVA with Tukey’s post-hoc test. * ≤0.05, ** ≤0.01, *** ≤0.001, **** ≤0.0001.

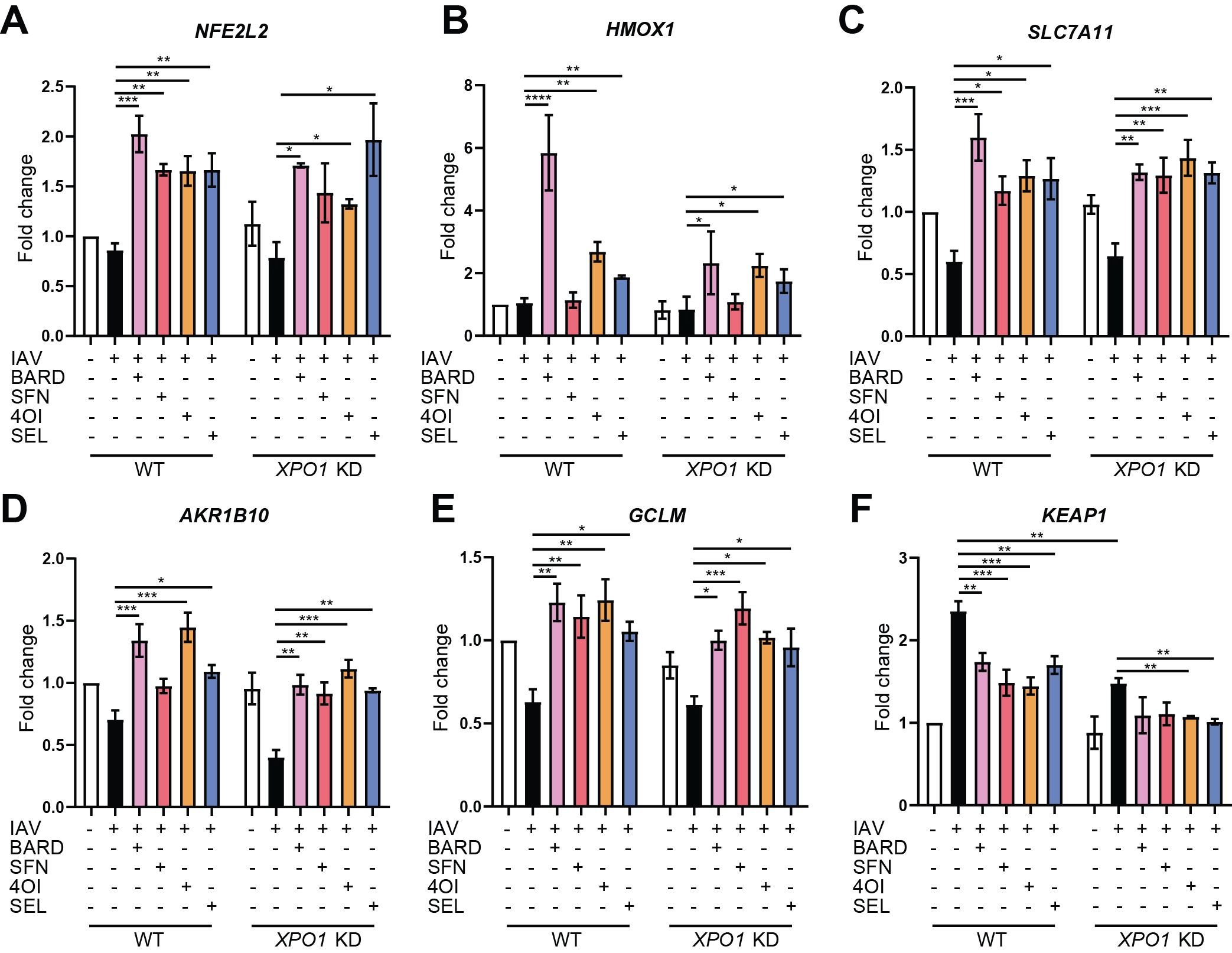

**Figure S2. Bar graphs corresponding to the heat map shown in Figure 3J.** The RT-qPCR data were analyzed by the 2^-ΔΔCt^ method using *HPRT* mRNA as internal control. Fold change was calculated with respect to expression in uninfected wild-type cells. *n*=3, means ±SEM. One-way ANOVA with Tukey’s post-hoc test. * ≤0.05, ** ≤0.01, *** ≤0.001, **** ≤0.0001.

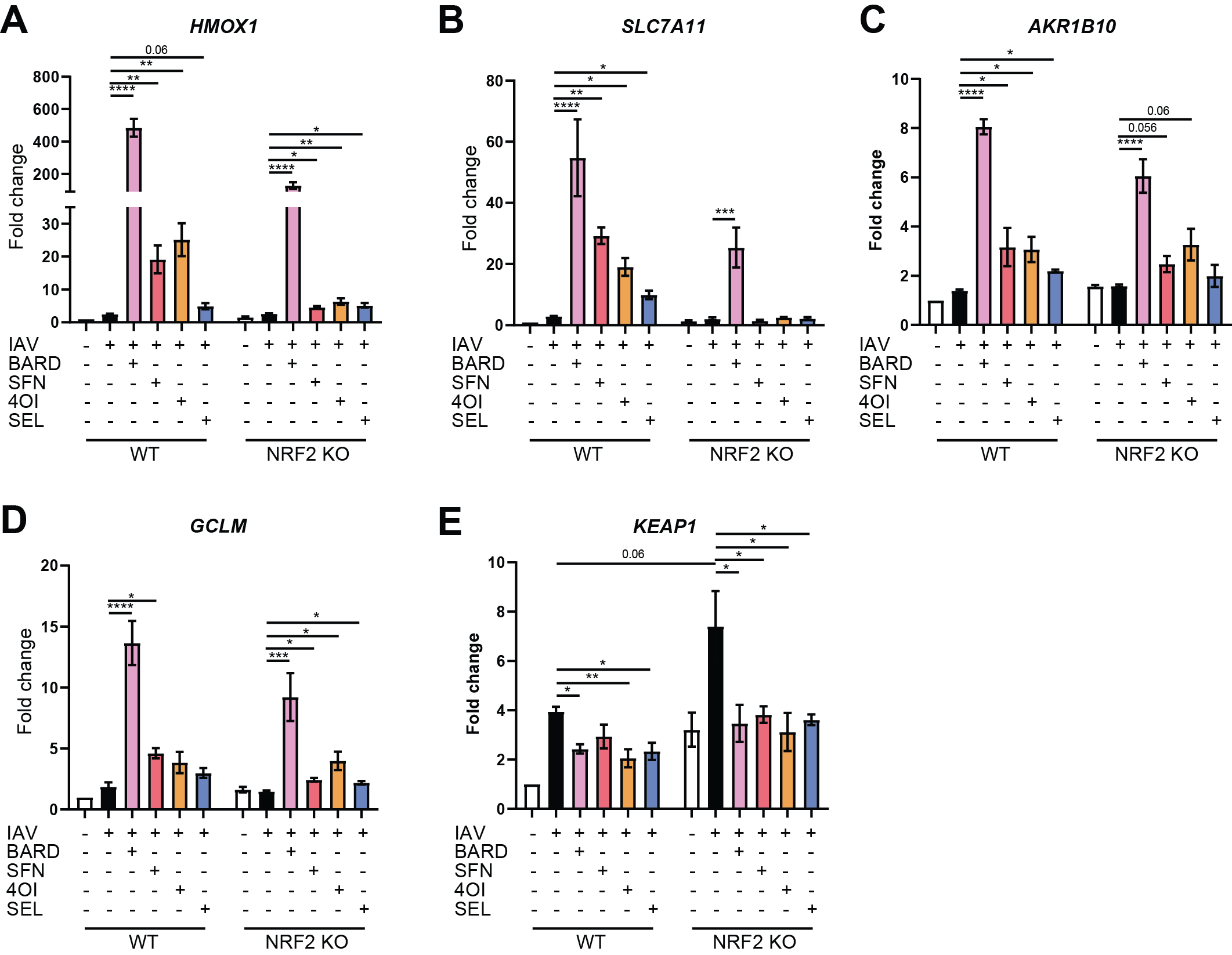

**Figure S3.** **Bar graphs corresponding to the heat map shown in Figure 4C.** The RT-qPCR data were analyzed by the 2^-ΔΔCt^ method using *HPRT* mRNA as internal control. Fold change was calculated with respect to expression in uninfected wild-type cells. *n*=3, means ±SEM. One-way ANOVA with Tukey’s post-hoc test. * ≤0.05, ** ≤0.01, *** ≤0.001, **** ≤0.0001.

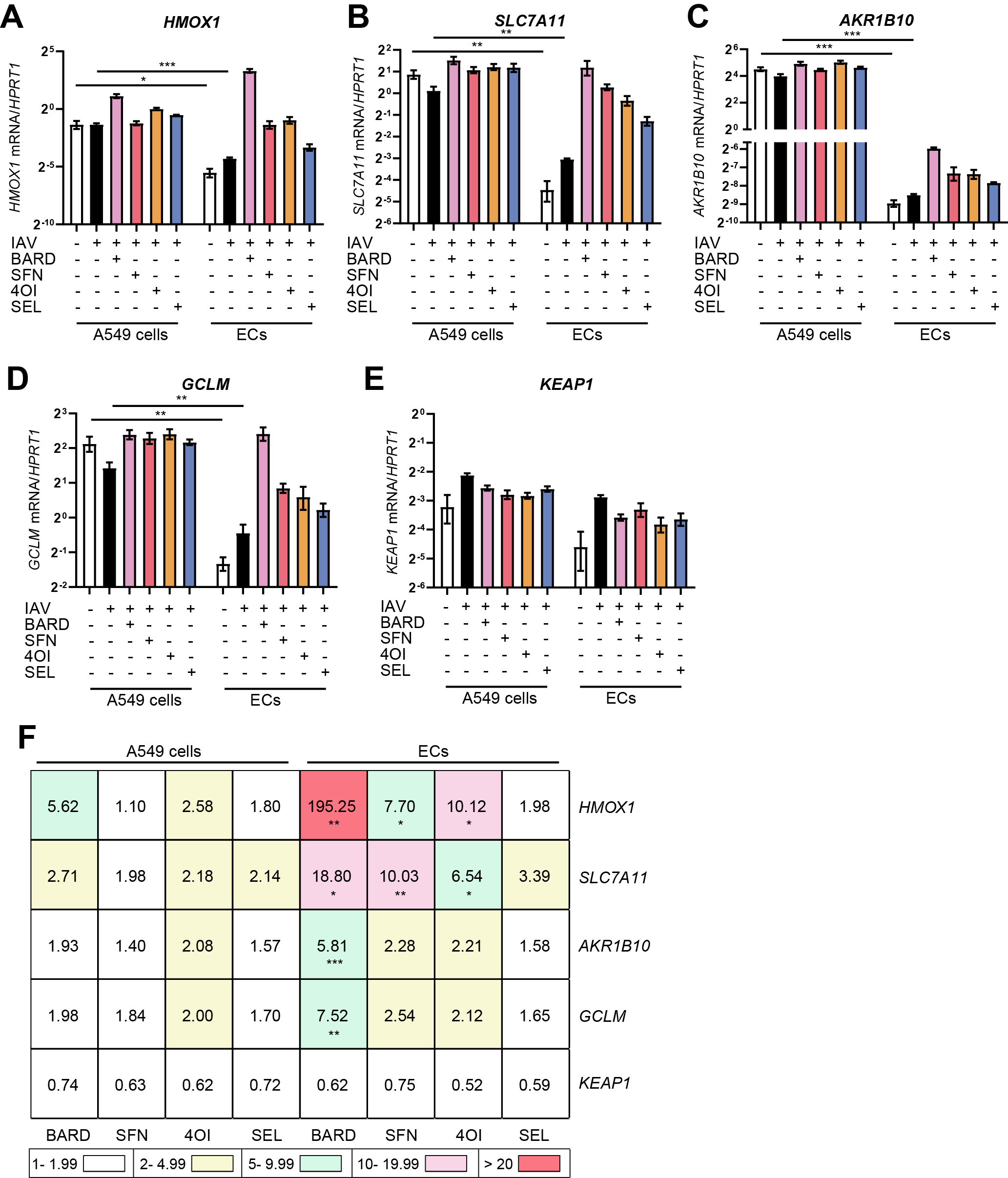

**Figure S4. The dynamic range of induction of anti-oxidative mRNAs by the four compounds is greater in iPSC-derived ECs than in A549 cells.** Reanalysis of the data of **Figure 3J** and **Figure 4C. A-D**, Baseline expression of the target genes is significantly higher in A549 cells than in iPSC-derived ECs. RT-qPCR data were reanalyzed using the 2^-ΔCt^ method, using *HPRT* mRNA as reference. Differences in expression (expressed on log_2_ scale) in the absence of treatment was compared between A549 and iPSC-derived ECs, either in uninfected or infected cells. *n*=3, means ±SEM. T-test. * ≤0.05, ** ≤0.01, *** ≤0.001, **** ≤0.0001. **E,** Induction of the target genes by the treatments is greater in iPSC-derived ECs than in A549 cells. Fold change (expressed as linear values) was computed for each cell type and compound by the 2^-ΔΔCt^ method, using infected untreated cells as reference. Differences between the two cell types in fold change due to the same treatment were assessed by T-test. * ≤0.05, ** ≤0.01, *** ≤0.001, **** ≤0.0001.

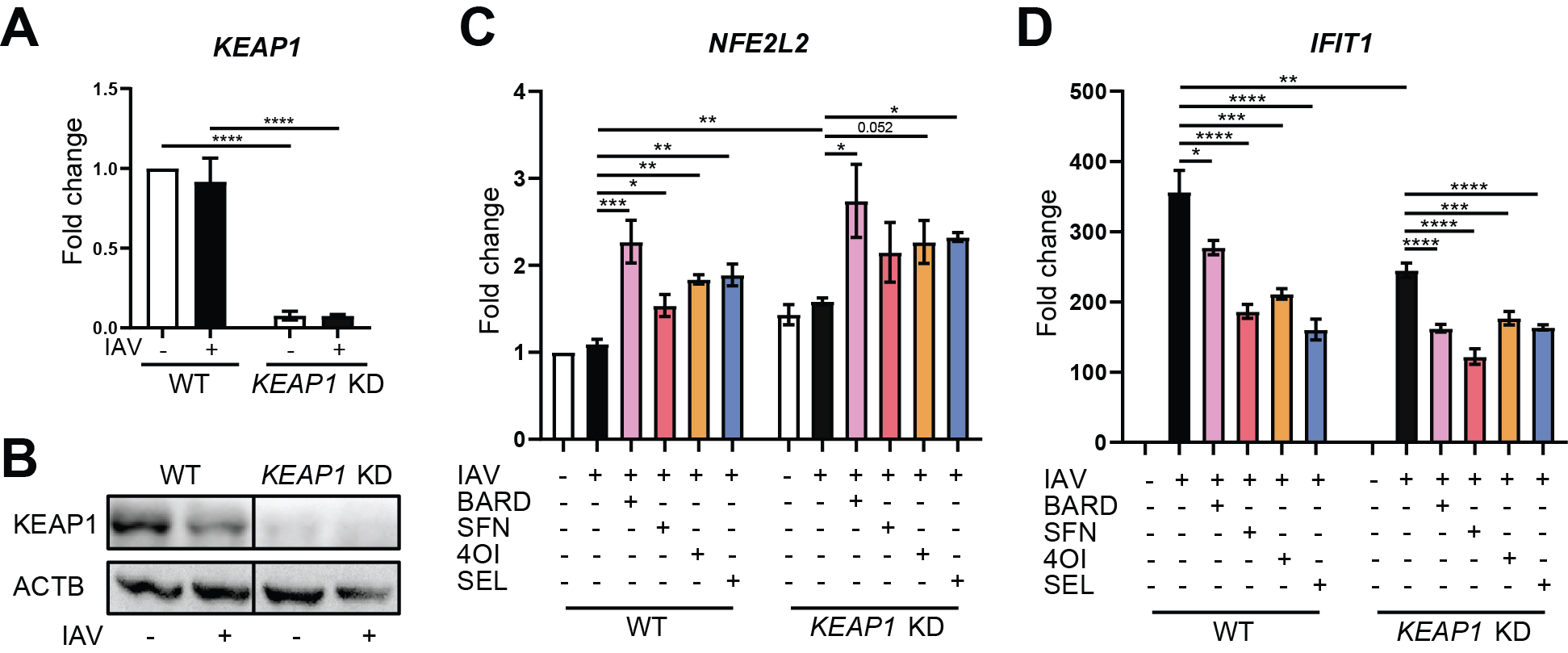

**Figure S5. Effects of KEAP1 knock-down on *NRF2 (NFE2L2)* and *IFIT1* mRNA expression.** A549 cells were transfected for 24 h with specific siRNA targeting *KEAP1* mRNA or nonspecific siRNA. Cells were then pretreated with the compounds (SEL, 1 µM; 4OI, 100 µM; BARD, 0.1 µM; SFN, 10 µM) for 12 h, infected with IAV PR8M (MOI=1) for 2 h, and then incubated in fresh buffer containing the compounds for 22 h. **A,B.** Efficiency of KEAP1 knock-down. **A.** *KEAP1* mRNA (RT-qPCR). **B.** KEAP1 protein (immunoblot). **C, D.** *NFE2L2* and *IFIT1* mRNA (RT-qPCR). n=3, means ±SEM. One-way ANOVA with Tukey’s post-hoc test, using infected untreated wild-type or knock-down cells as reference. * ≤0.05, ** ≤0.01, *** ≤0.001, **** ≤0.0001.

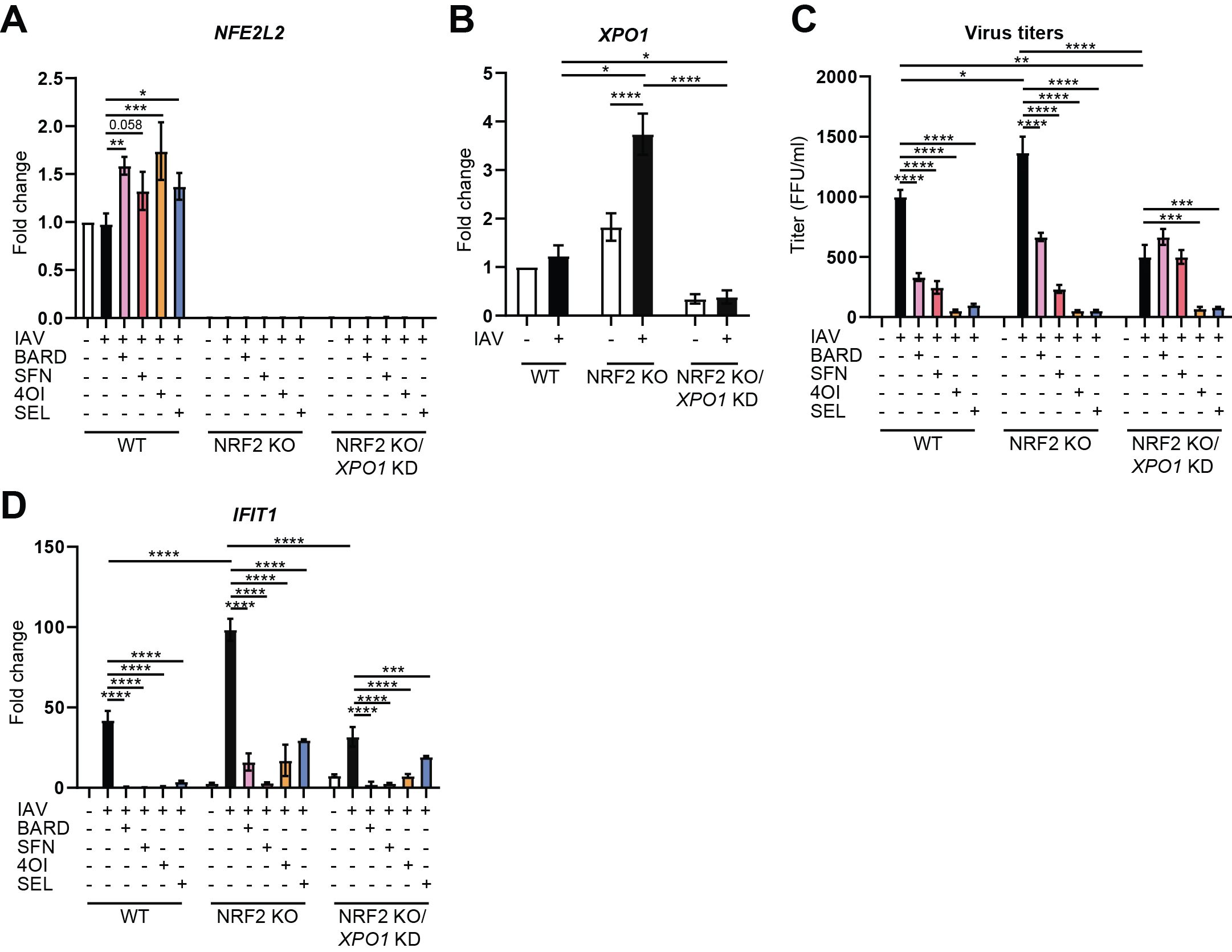

**Figure S6. Effects of *XPO1* knock-down in *NRF2^-/-^* ECs on antiviral activity of the compounds.** WT and *NRF2^-/-^* ECs were transfected for 24 h with specific siRNA targeting *XPO1* mRNA or nonspecific siRNA. Cells were then pretreated with the compounds (SEL, 1 µM; 4OI, 100 µM; BARD, 0.1 µM; SFN, 10 µM) for 12 h, infected with IAV PR8M (MOI=1) for 2 h, and then incubated in fresh buffer containing the compounds for 22 h. **A.** *NRF2* *(NFE2L2)* mRNA (RT-qPCR). **B.** *XPO1* mRNA (RT-qPCR), demonstrating a 75% knock-down. **C.** IAV titers in cell culture supernatants (foci-forming assay, FFU/mL). **D.** *IFIT1* mRNA (RT-qPCR). n=3, means ±SEM. One-way ANOVA with Tukey’s post-hoc test, using infected untreated wild-type or knock-down cells as reference. * ≤0.05, ** ≤0.01, *** ≤0.001, **** ≤0.0001.

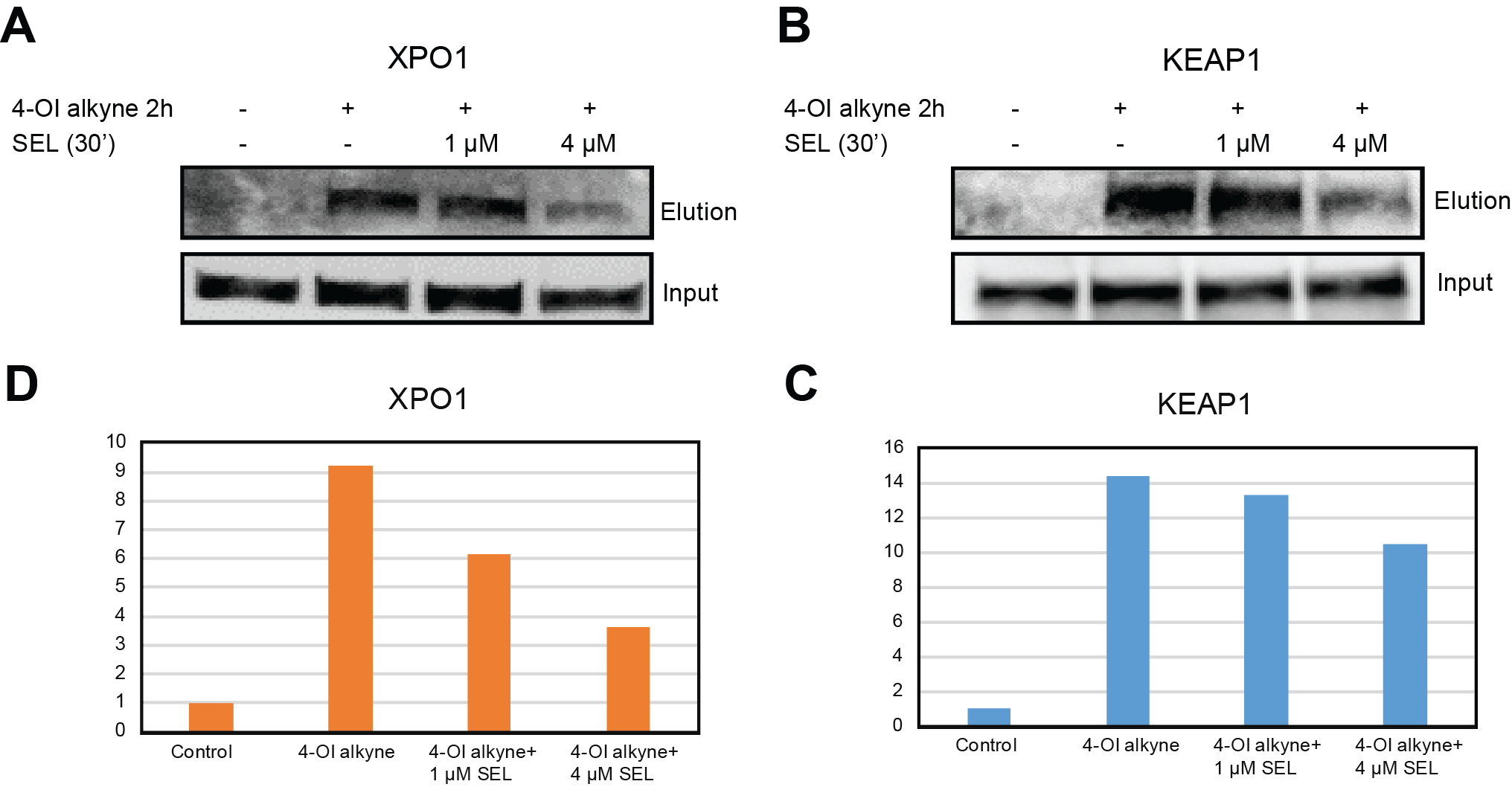

**Figure S7. Competition experiment demonstrating binding of 4-OI and SEL to the same sites on XPO1 and KEAP1. A,B.** “Click-chemistry” pull-down assay demonstrating covalent binding of an alkynated 4OI probe (4-OI-alk) to XPO1 (A) and KEAP1 (B) in Calu-3 cells. Cells were preincubated with 1 or 4 µM unmodified SEL for 30 min. as indicated. Two hours after addition of the probe, proteins complexed with the probe were detected by immunoblot for XPO1 (A) or KEAP1 (B). **C,D.** Densitometry (arbitrary units) of the immunoblots, normalized to the signal obtained from the band labeled “input”. SEL competes with 4OI for complex formation with both targets, suggesting that the compounds recognize the same sites on both targets. However, competition is less efficient for complex formation with KEAP1, suggesting that 4OI has higher affinity for KEAP1 and that SEL has higher affinity for XPO1.

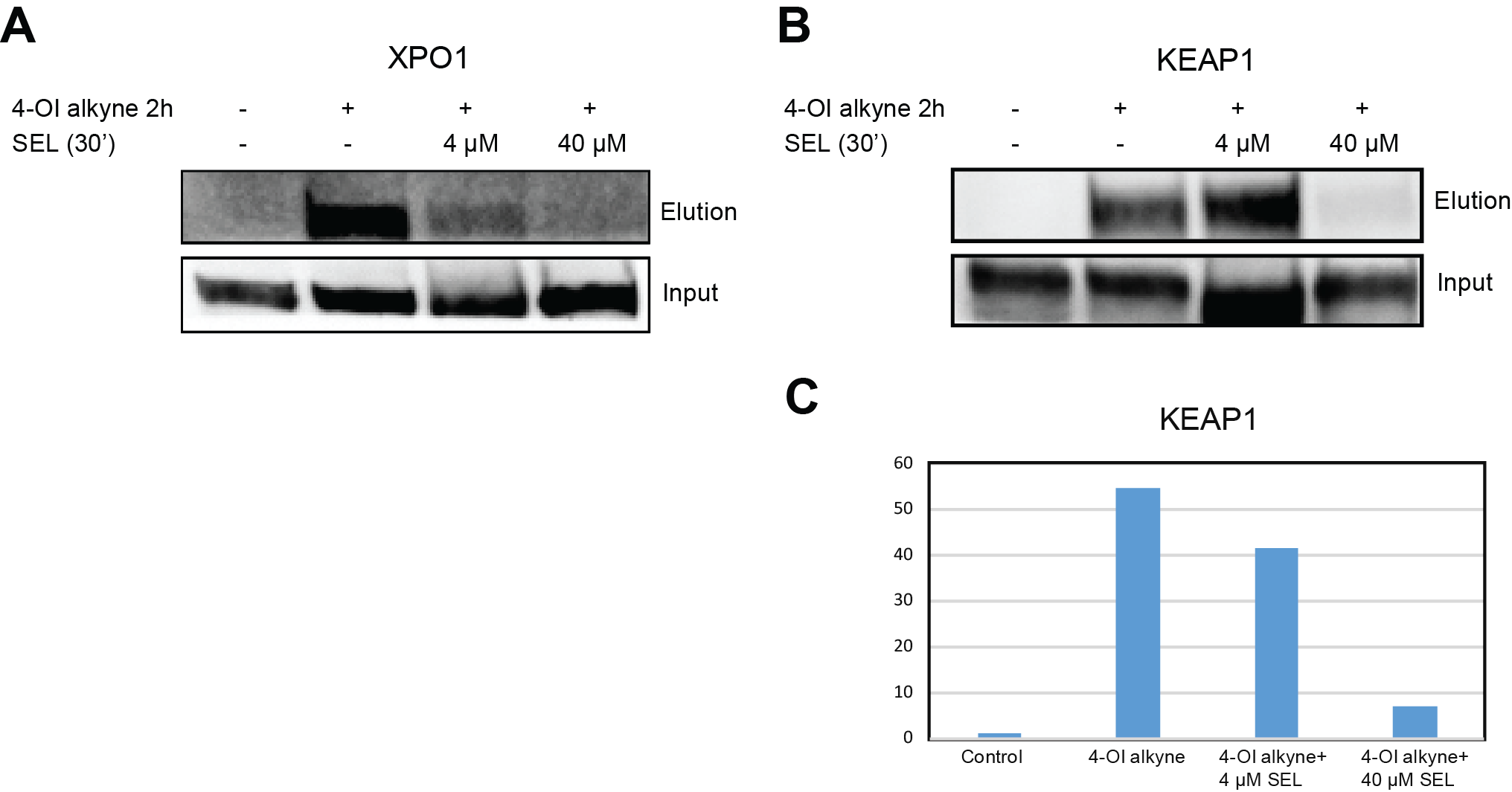

**Figure S8. Experiment identical as Figure S6, but featuring a higher concentration of SEL.** SEL was added at concentrations of 4 and 40 µM. Due to a technical error, the signals in **B** (4 µM SEL) were higher than expected, but densitometry revealed a similar reduction in complex formation as in the experiment shown in Figure S6. Densitometry could not be performed on **A** due to loss of part of the membrane (see missing lower border of band “Input 4 µM”).

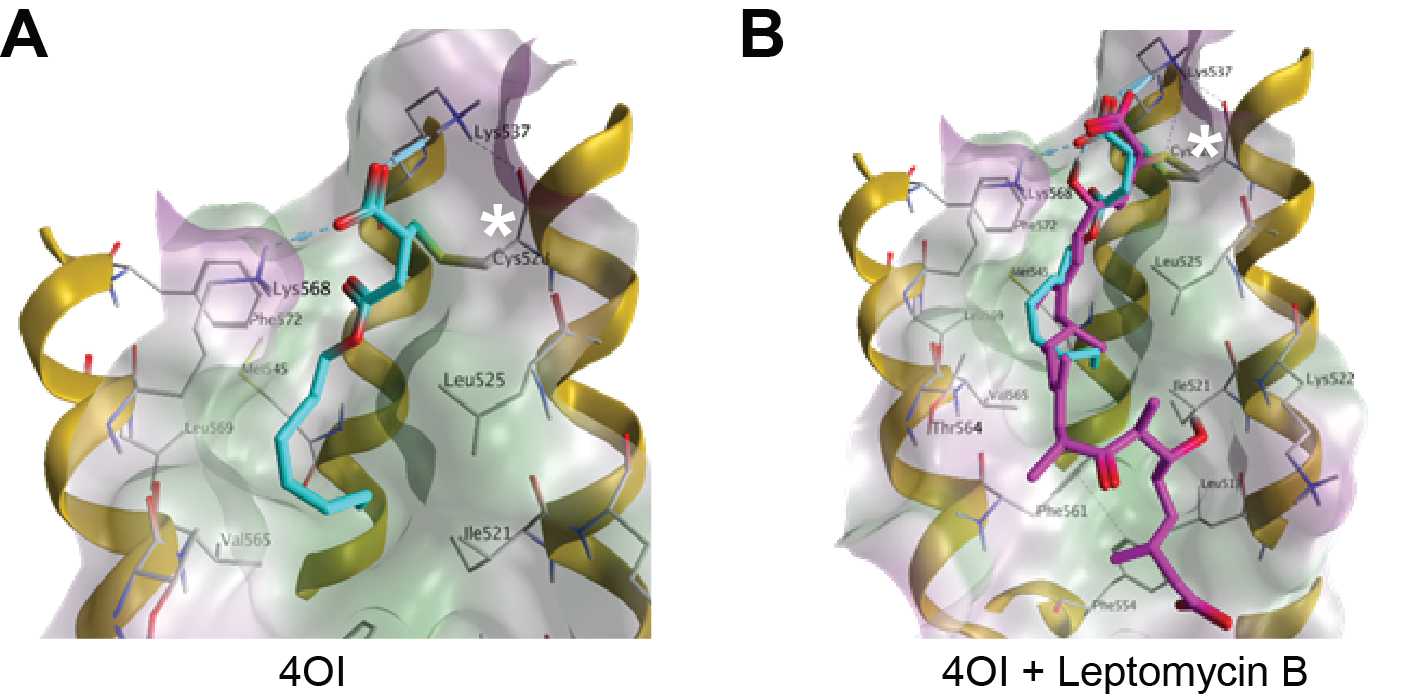

**Figure S9.** **3D structural modeling of 4OI-XPO1 interactions based on the co-crystal structure of XPO1 (CRM1) with leptomycin B (PDB ID: 6TVO)**. Both 4OI and leptomycin B are covalently bound to the reactive Cys528 (marked with an asterisk *) and interact extensively with the hydrophobic NES-binding groove. **A.** 4OI binds the site through hydrophobic interactions between the octyl chain and Ile521, Leu525, Met545, Val565 and Leu569 in the hydrophobic pockets Φ2 and Φ3 of the NES-binding site. The C1-carboxyl group further stabilizes binding through two hydrogen bonds with Lys537 and Lys568. These hydrophobic and electrostatic interactions optimally direct the methylene group of 4OI towards Cys528 and could be the driving force for the covalent Michael 1,4-addition. **B.** Overlay of 4OI (cyan) and leptomycin B (magenta) in the NES-binding groove showing about 70% occupancy by leptomycin B and 40% by 4OI. Lipophilicity protein surface at the NES-binding cleft: lipophilic (green), hydrophilic (violet), neutral (white), α-helices (gold). * = Cys528.

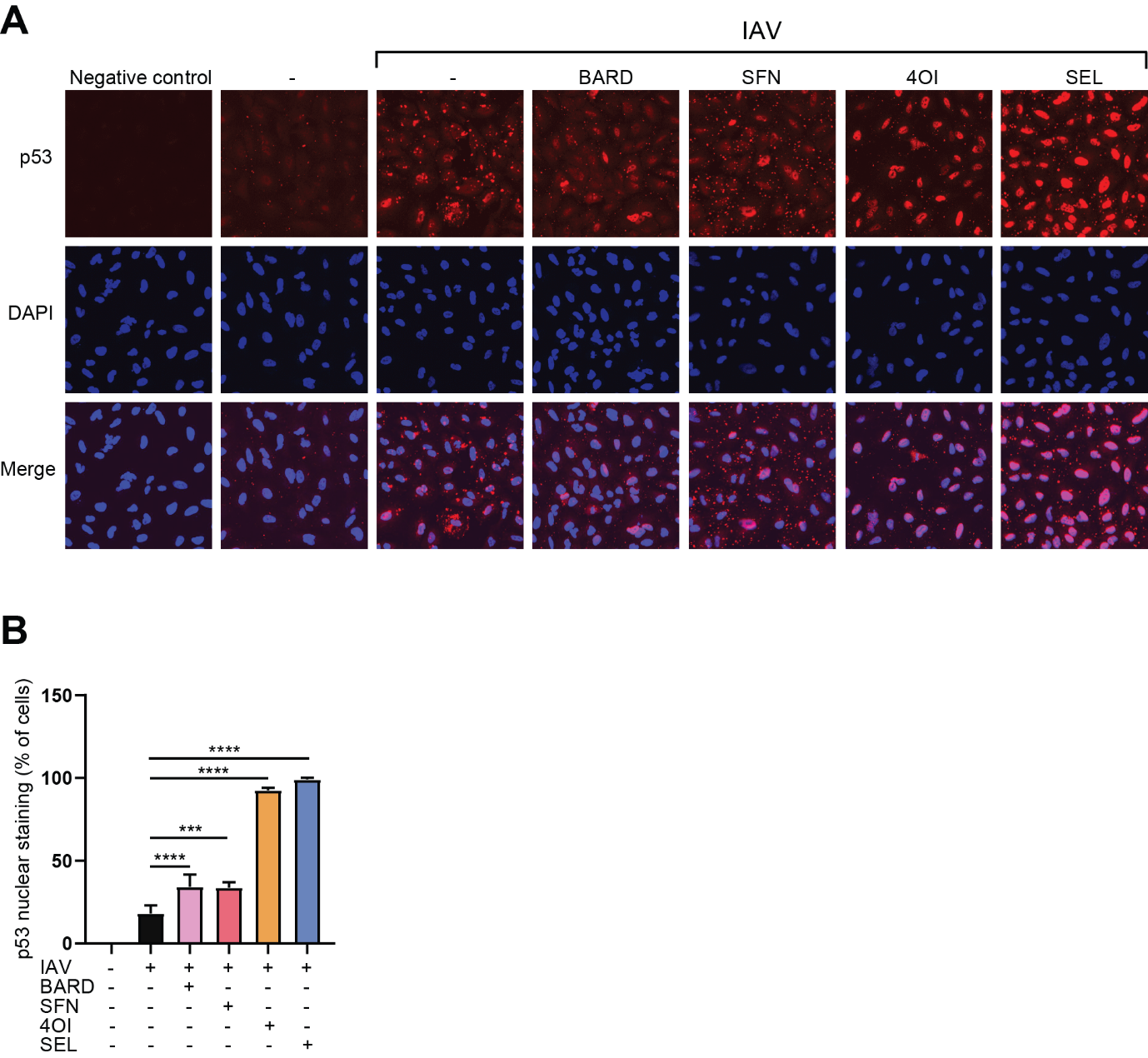

**Figure S10.** **The compounds favor nuclear retention of p53 in IAV-infected A549 cells.** A549 cells were treated and infected as described for Figure 2. p53 was detected by indirect immunofluorescence 8 h p.i., using Alexa Fluor 568 labeled secondary antibody. **A.** Representative immunofluorescence images p53 = red. Nuclei = blue (DAPI). Pink signal in merged images = nuclear localized p53. The positive staining granular pattern is a technical artefact and was considered background signal. Negative control = no primary antibody. **B.** Fraction of all cells with nuclear p53 staining. Cells with nuclear p53 staining were counted by visual inspection by two independent examiners who were blinded to the identity of the specimens. n=4 microscopic fields, means ±SEM. One-way ANOVA with Tukey’s post-hoc test, using infected untreated wild-type or knock-down cells as reference. * ≤0.05, ** ≤0.01, *** ≤0.001, **** ≤0.0001.

| **Table S1.** Viruses with pharmacologic evidence of XPO1 (CRM1)-dependence | | |
| --- | --- | --- |
| **Virus** | **Family** | **References** |
| IAV  SARS-CoV-2  HIV-1  HTLV-1  RSV  DENV  RV  HCMV | *Orthomyxoviridae*  *Coronaviridae*  *Retroviridae*  *Retroviridae*  *Pneumoviridae*  *Flaviviridae*  *Rhabdoviridae*  *Herpesviridae* | 1, 2, 3, 4, 5  6  4  4  4, 5  4  4  4 |
| **Abbreviations**  Dengue Virus (DENV), Human Immune Deficiency Virus Type 1 (HIV-1), Human T-Cell Leukemia Virus type 1 (HTLV-1), Human Cytomegalo Virus (HCMV), Influenza A Virus (IV), Respiratory Syncytial Virus (RSV), Rabies Virus (RV), Severe Acute Respiratory Syndrome Coronavirus (SARS-CoV-2). | | |
| **References**  1. Elton D et al. **Interaction of the Influenza Virus Nucleoprotein with the Cellular CRM1-Mediated Nuclear Export Pathway.** Journal of Virology Volume 75, Issue 1, 1 January 2001, Pages 408-419. DOI: <https://doi.org/10.1128/JVI.75.1.408-419.2001>  2. Chutiwitoonchai N et al. **Inhibition of CRM1-mediated nuclear export of influenza A nucleoprotein and nuclear export protein as a novel target for antiviral drug development**. Volume 507, July 2017, Pages 32-39. DOI: <http://dx.doi.org/10.1016/j.virol.2017.04.001>  3. Chutiwitoonchai N and Aida Y. **NXT1, a Novel Influenza A NP Binding Protein, Promotes the Nuclear Export of NP via a CRM1-Dependent Pathway**. *Viruses* 2016, *8*(8), 209; DOI: [**https://doi.org/10.3390/v8080209**](https://doi.org/10.3390/v8080209)  4. Mathew C and Ghildyal R. **CRM1 Inhibitors for Antiviral Therapy.** *Front. Microbiol*., 28 June 2017. Sec. Pickens Antimicrobials, Resistance and Chemotherapy. Volume 8 – 2017. DOI: **<https://doi.org/10.3389/fmicb.2017.01171>**  5. Pickens JA and Tripp RA. **Verdinexor Targeting of CRM1 is a Promising Therapeutic Approach against RSV and Influenza Viruses**. Viruses 2018, 10(1), 48. DOI: <https://doi.org/10.3390/v10010048>)  6. Kashyap T, Murray J, Walker CJ, Chang H, …, Ripp RA, Landesman Y. Selinexor, a novel selective inhibitor of nuclear export, reduces SARS-CoV-2 infection and protects the respiratory system in vivo. *Antiviral Res*. 2021, 192:105115. ddoi: 10.1016/j.antiviral.2021 | | |

| **Table S2.** List of RT-qPCR primers | | |
| --- | --- | --- |
| **Gene name** | **Primer name** | **Sequence (5´ - 3´)** |
| ***IFIT1*** | IFIT1-F | TCAGGCATTTCATCGTCATC |
|  | IFIT1-R | GCAGAACGGCTGCCTAATTT |
| ***CXCL10*** | CXCL10-F | CTGCTTTGGGGTTTATCAGA |
|  | CXCL10-R | CCACTGAAAGAATTTGGGC |
| ***HMOX1*** | HMOX1-F | AGGGAAGCCCCCACTCAAC |
|  | HMOX1-R | ACTGTCGCCACCAGAAAGCT |
| ***NFE2L2*** | NFE2L2-F | CAGCGACGGAAAGAGTATGA |
|  | NFE2L2-R | TGGGCAACCTGGGAGTAG |
| ***HPRT1*** | HPRT-F | GAACGTCTTGCTCGAGATGTG |
|  | HPRT-R | CCAGCAGGTCAGCAAAGAATT |
| ***HA*** | HA-F | CTCGTGCTATGGGGCATTCA |
|  | HA-R | TTCCAATCGTGGACTGGTGT |
| ***XPO1*** | XPO1-F | TGGGCTGAAAACTCAACCGAG |
|  | XPO1-R | TTGCTGATGCTGTAGCTCCC |
| ***SLC7A11*** | SLC7A11-F | TCCTGCTTTGGCTCCATGAACG |
|  | SLC7A11-R | AGAGGAGTGTGCTTGCGGACAT |
| ***AKR1B10*** | AKR1B10-F | GAGGACCTGTTCATCGTCAGCA |
|  | AKR1B10-R | CGTCCAGATAGCTCAGCTTCAG |
| ***GCLM*** | GCLM-F | TCTTGCCTCCTGCTGTGTGATG |
|  | GCLM-R | TTGGAAACTTGCTTCAGAAAGCAG |
| ***KEAP1*** | KEAP1-F | TTCGCCTACACGGCCTC |
|  | KEAP1-R | GAAGTTGGCGATGCCGATG |

### **Supplemental Methods**

### **Organic synthesis – general methods**

All reactions were conducted in flame-dried glassware under an atmosphere of dry argon unless otherwise stated. CH_2_Cl_2_ was dried over activated 4Å molecular sieves and reagents were used as received from commercial suppliers (Sigma Aldrich, TCI and Fluorochem). Concentration in vacuo was performed using a rotary evaporator with the water bath temperature at 40 °C, followed by further concentration using a high vacuum pump. TLC analysis was carried out on silica coated aluminum foil plates (Merck Kieselgel 60 F254). The TLC plates were visualized by UV irradiation and/or by staining with KMnO4 stain. Purification was performed by automated flash column chromatography (AFCC) using an Interchim PuriFlash 420 instrument with 30 µm prepacked columns. Infrared spectra (IR) were acquired on a PerkinElmer Spectrum TwoTM UATR. Mass spectra (HRMS) were recorded on a Bruker Daltonics MicrOTOF time-of-flight spectrometer. Nuclear magnetic resonance (NMR) spectra were recorded on a Bruker BioSpin GmbH 400 MHz spectrometer, running at 400 and 101 MHz for ^1^H and ^13^C, respectively. The residual peak of the respective solvent was used as the internal standard: DMSO-*d*_6_ (CD_2_HSOCD_3_ *δ*H 2.50 ppm, CD_3_SOCD_3_ *δ*C 39.5 ppm).

Synthesis of *2-Methylene-4-(oct-7-yn-1-yloxy)-4-oxobutanoic acid* (**4-OI-alk**)

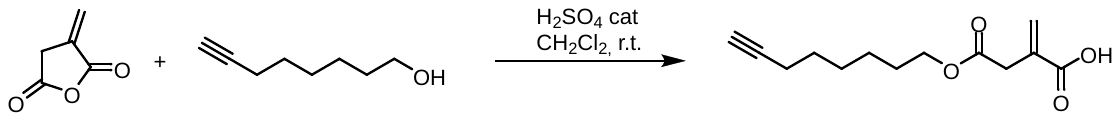

Itaconic anhydride (100 mg, 0.89 mmol, 1.0 eq.) was dissolved in anhydrous CH_2_Cl_2_ (0.5 mL). 7-Octyn-1-ol (338 mg, 2.67 mmol, 3.0 eq.) and conc. H_2_SO_4_ (5 drops) were added, and the reaction mixture was stirred for 16 hours at r.t. Et_2_O (25 mL) was then added to the reaction mixture and the organic phase was washed with an aqueous solution of K_2_CO_3_ (10 w%, 2 × 10 mL). The aqueous phase was then extracted with Et_2_O (2 × 20 mL) to remove the non-ionizable impurities. Conc. HCl was then added to the aqueous phase until pH = 1 and
the aqueous phase was then extracted with CH_2_Cl_2_ (2 × 25 mL). The combined organic phases were dried over Na_2_SO_4_, filtered, and then concentrated under reduced pressure. The crude product was purified by flash chromatography on a silica gel column using 10-100% EtOAc in heptane as eluent to give **4-OI-alk** (70 mg, 0.30 mmol, 33%) as a white solid.

R*_f_ =* 0.50 (Pentane/EtOAc 2:1; UV (254 nm) and KMnO_4_

HRMS (ESI) m/z calcd for C_13_H_17_O_4_ [M-H]^-^ 237.1132, found 237.1131

IR ν_max_ (cm^-1^) 3294, 2939, 1735, 1698, 1634, 1432, 1160, 960, 635 νννν

^1^H NMR (400 MHz, DMSO-*d*_6_) *δ* 12.62 (s, 1H), 6.15 (d, *J* = 1.6 Hz, 1H), 5.76 (d, *J* = 1.5 Hz, 1H), 4.00 (t, *J* = 6.6 Hz, 2H), 3.30 (s, 2H), 2.75 (t, *J* = 2.7 Hz, 1H), 2.15 (td, *J* = 6.9, 2.7 Hz, 2H), 1.58 – 1.49 (m, 2H), 1.46 – 1.39 (m, 2H), 1.43 – 1.22 (m, 4H).

^13^C NMR (101 MHz, DMSO-*d*_6_) *δ* 171.0, 167.8, 135.3, 128.4, 85.0, 71.7, 64.5, 37.7, 28.4, 28.3, 28.2, 25.2, 18.1.

*The NMR spectra are in agreement with the values reported in ref. 18 (main text).*

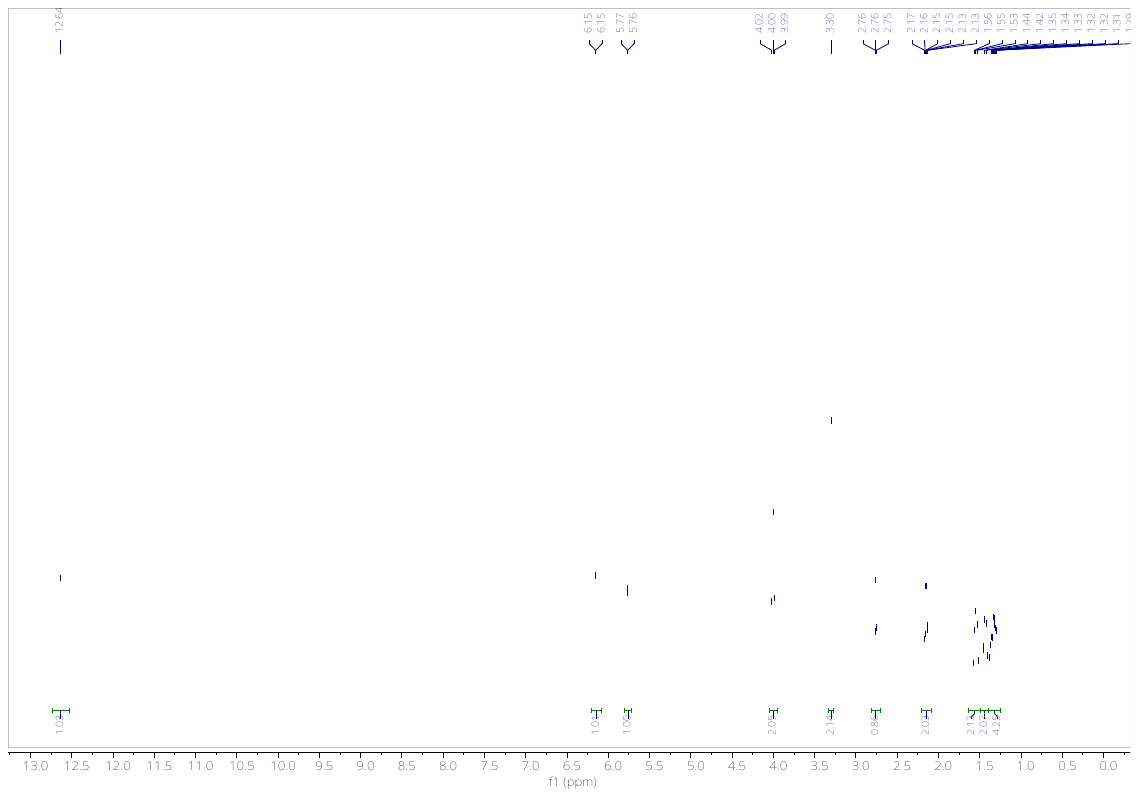

**Figure S11:** ^1^H NMR spectrum (400 MHz, DMSO-*d*_6_) of **4-OI-alk**

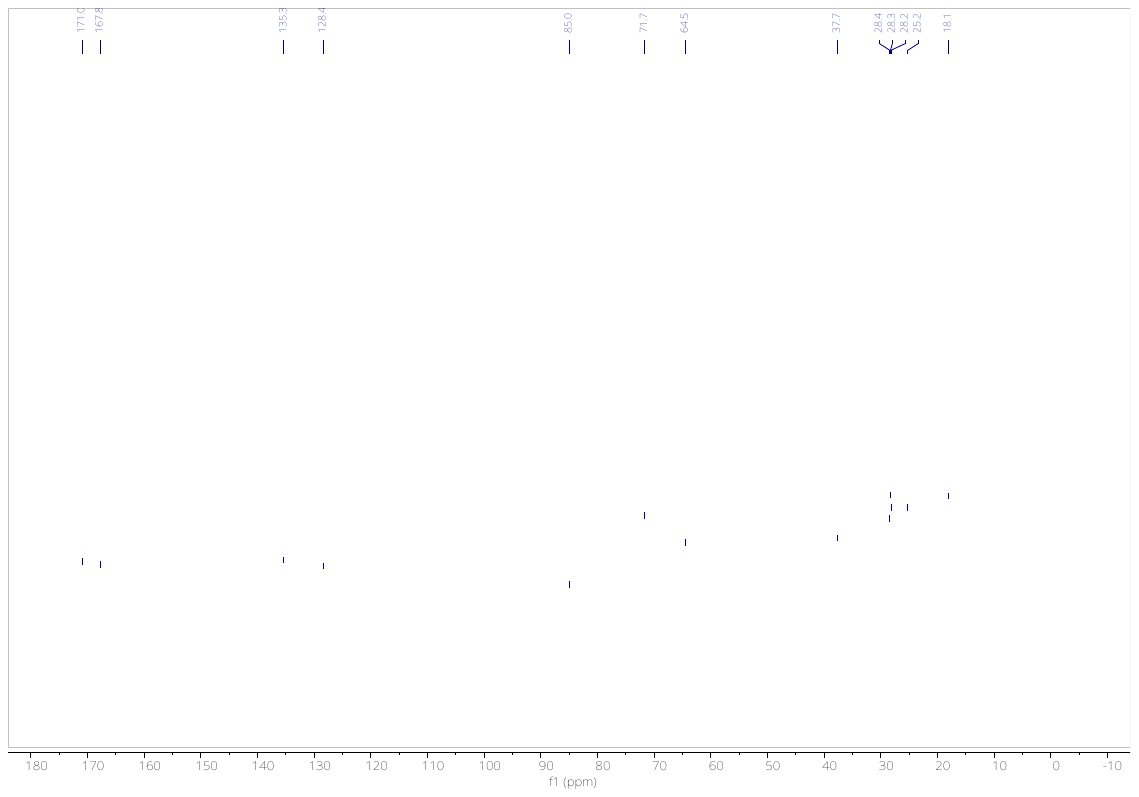

**Figure S12:** ^13^C NMR spectrum (101 MHz, DMSO-*d*_6_) of **4-OI-alk**

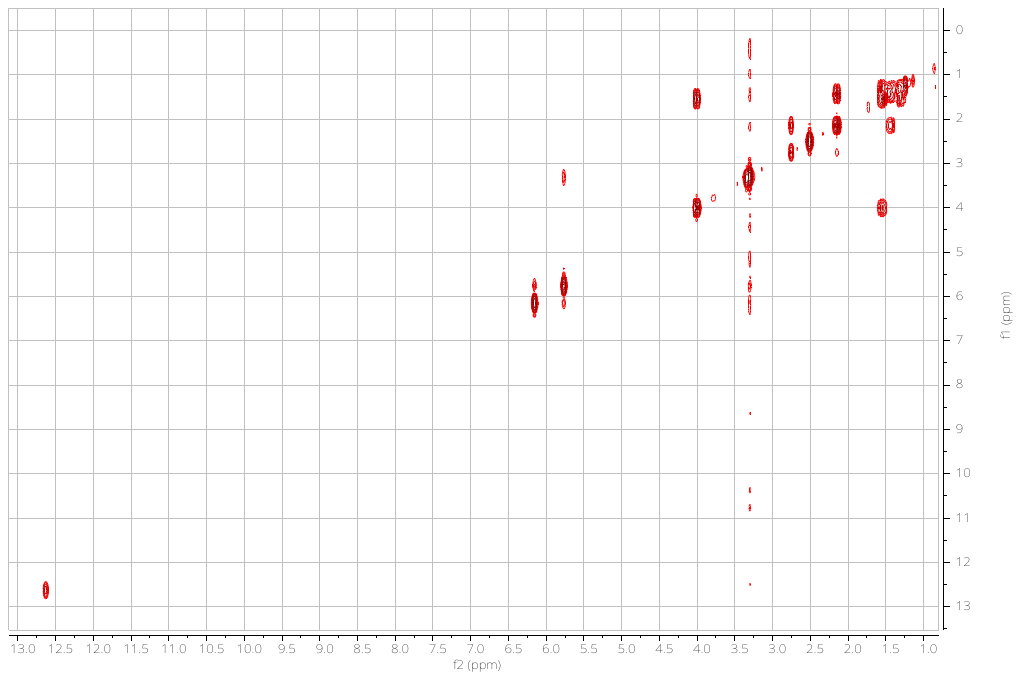

**Figure S13**: COSY spectrum (400 MHz, DMSO-*d*_6_) of **4-OI-alk**

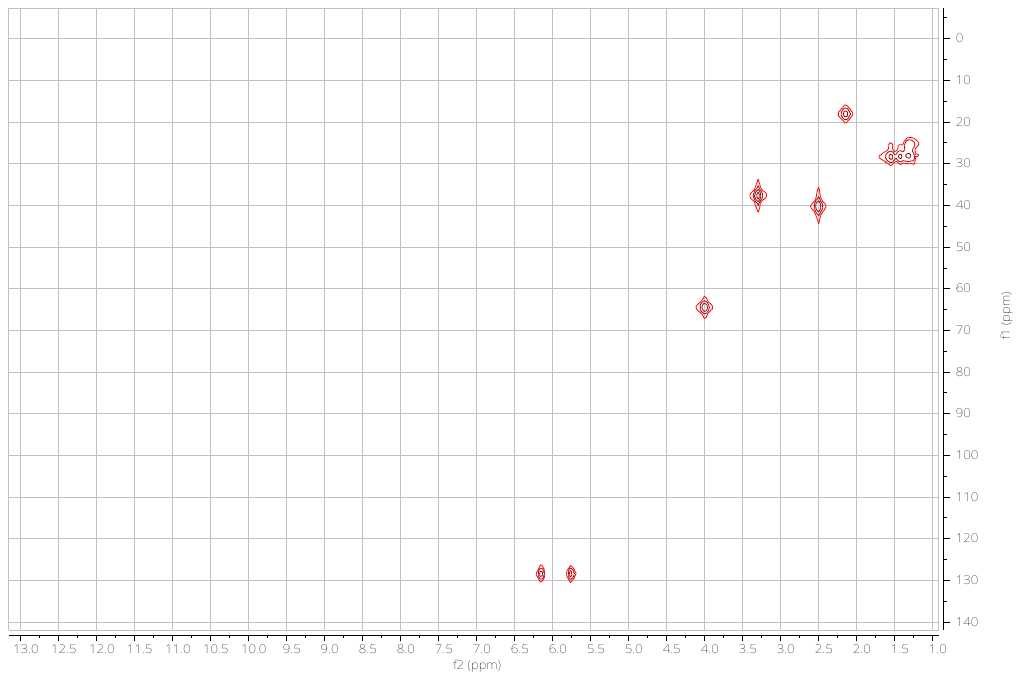

**Figure S14**: HSQC NMR spectrum (400/101 MHz, DMSO-*d*_6_) of **4-OI-alk**

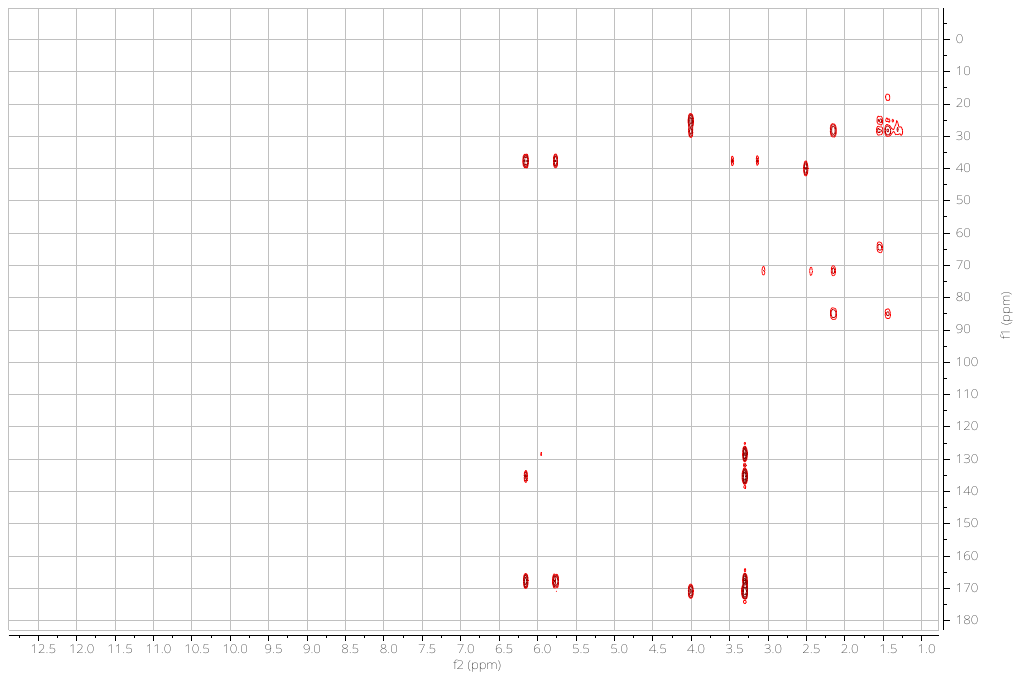

**Figure S15**: HMBC NMR spectrum (400/101 MHz, DMSO-*d*_6_) of **4-OI-alk**
